# Assessment of the bovine uterus with endometritis using Doppler ultrasound

**DOI:** 10.1101/826065

**Authors:** Bruno Leonardo Mendonça Ribeiro, Enoch Brandão de Souza Meira Júnior, Mario Augusto Aleman Reyes, Eduardo Carvalho Marques, Alessandra Figueiredo de Castro Nassar, Lilian Gregory

## Abstract

Postpartum uterine diseases such as metritis and endometritis are highly prevalent in dairy cows. These diseases negatively affect the reproductive performance and consequently economic activity. Cows in the puerperal period presenting endometritis may have alterations in the hemodynamics of the uterine tissue and the uterine arteries, which differ them from healthy cows. Therefore, this study aimed to use the Doppler ultrasonography to describe the hemodynamic changes in the uterus of cows showing endometritis diagnosed between 25 - 35 days postpartum. Eighty-nine Holstein Friesian females with 25 to 35 days postpartum were studied. Cows were assigned to two experimental groups, infected or not infected, according to the results of the endometrial cytology. Clinical examination, vaginoscopy, Doppler ultrasound and sample collection (saline solution was injected and recovered by endoscopie method aiming cytological and microbiological evaluation of the uterus) were also performed. Cows with endometritis had the cervix (*P* = 0.040) and the left horn (*P* = 0.020) increased compared to healthy cows. 78.6% of the endometritic cows showed abnormal uterine discharge, while 57.6% of healthy cows had this same condition (*P =* 0.0005). The spectral Doppler evaluation of the uterine arteries revealed no differences between groups. *Bacillus spp., Trueperella pyogenes, Escherichia coli* and *Staphylococcus intermedius* were the most isolated bacteria among samples. Higher score or increase of uterus vascularization of the endometrial Doppler was correlated with *Trueperella pyogenes* (*P =* 0.0003) and intrauterine heterogeneous content (*P* = 0.0047). Finally, mesometrial Doppler was correlated with endometrial Doppler (*P* < 0.001), uterine bacteria (*P =* 0.001) and intrauterine heterogeneous content (*P =* 0.049). Regarding the evolution of uterine alterations, Doppler ultrasonography provides fast results and is a lesser invasive technique such as uterus biopsie and endometrial citology and gives answers about fertility and uterus health.

## 1. INTRODUCTION

High milk production causes high negative energy balance and could cause changes in hormone levels, embryonic losses and a higher incidence of reproductive disorders in dairy cows [1,2]. Unsatisfactory reproductive performance negatively affects productivity [3], decreasing milk production and number of calves [4].

The uterine infection, like metritis and endometritis is the most important cause of infertility in dairy cows. Therefore causing decrease in reproductive performance with economic injuries [5,6]. Metritis is defined as a severe alteration of all layers of the uterus. In the clinical uterine examination the infected has reddish-brown fetid uterine discharge and systemic clinical signs such as fever and reduced milk production, normally within 21 days postpartum [7]. In contrast endometritis occurs after 21 days postpartum and is the inflammation of the endometrium. Animals show no systemic clinical signs [8]. Cytological endometritis may also occur, with absence of purulent discharge but high number of neutrophils on the endometrial cytological slide[8].

By showing the presence, direction and type of blood flow the Doppler ultrasonography technique determines hemodynamics changes wich are important to understand the morphophysiological aspects of the female reproductive tract [9]. This concept is used in women ginecology to tell a part fertile [10] and infertile women [11]. Doppler ultrasound shows up the changes in the uterine circulation and it is used widely in veterinary reproduction [12,13].

It is hypothesized that alterations in the hemodynamics of the uterus tissue and arteries may happened in cows presenting endometritis in the puerperal. Thus, the use of more than one diagnostic technique may help in the identification of uterine diseases. Therefore, this study aimed to use the Doppler ultrasonography to describe the hemodynamic changes in the uterus of cows showing endometritis diagnosed between 25 - 35 days postpartum.

## 2. MATERIALS AND METHODS

### 2.1 Area characterization and case definition

This study was carried out in dairy farms from the State of São Paulo (N = 2) and from the State of Minas Gerais (N=1), Brazil, under the strict regulations of CONCEA (Brazilian commetee for Animal use in experiments, protocol number 5135030214 and approved 27/08/2014 at CEUA, commetee for animal use in experiments at the Veterinary School in the University of São Paulo), equivalent to EU Directive 2010/63/EU for animal experiments. All farms have freestall breeding system and assisted calving. Eighty-nine Holstein Friesian females with 25 to 35 days postpartum were studied. Cows were assigned to two experimental groups according to the results of the endometrial cytology [14] as follows:

a. Control group (C): healthy cows (neutrophil polymorphonuclear count < 10%).
b. Endometritis group (E): cows diagnosed by cytology (neutrophil polymorphonuclear count ≥10%).

Clinical examination, vaginoscopy, Doppler ultrasound and sample collection were also performed, all in the same day.

### 2.2 Clinical examination

Female genital tract was evaluated by rectal palpation as proposed by Grunert, Birgel and Vale (2005) [15]. Briefly, ovaries were evaluated searching for follicles and corpus luteum (CL). In the cervix, morphological alterations such as volume was evaluated. In the uterus, location, size, symmetry of the horns and consistency were also investigated. Changes were described according to Grunert, Birgel and Vale (2005) [15] using the classification: from I (not very thick) to VI (very bulky). The horns were classified as symmetrical (s) and asymmetrical (As). According to consistency (C), the uterus was classified as flaccid (CI), reactive (CII), and with vigorous and prolonged contraction (CIII).

### 2.3 Vaginoscopy

Vaginoscopy was performed after rectal palpation. The speculum was introduced in the vagina to observe the vaginal mucosa, cervix and the presence of discharge according to Grunert, Birgel and Vale (2005) [15].

### 2.4 Ultrasonography and Doppler mode

The ultrasound was performed with a m-turbo, Fuji film Sonosite (Bothel, WA, USA) with a micro convex probe (8-5 mhz), as proposed by Meira Jr et al. (2012) [16] and Heppelmann, Krüger and Leidl (2013)[17]. Diameters of the cervix and horns, ovarian structures and the presence of uterine discharge were evaluated. Cervical and uterine diameters were classified as small or negative (< 3.5 cm), medium or suspect (between 3.5 and 5 cm) and large or positive (> 5 cm) (Figure 1). Uterine dimensions were evaluated according to Meira Jr et al. 2012 [16] (Figure 1). The presence of heterogeneous intrauterine contents (hic) and the uterine wall with hyperechoic characteristics were associated with purulent discharge according to Descoteaux, Gnemmi and Colloton (2010) [18] (Figure 2).

**Figure 1:**
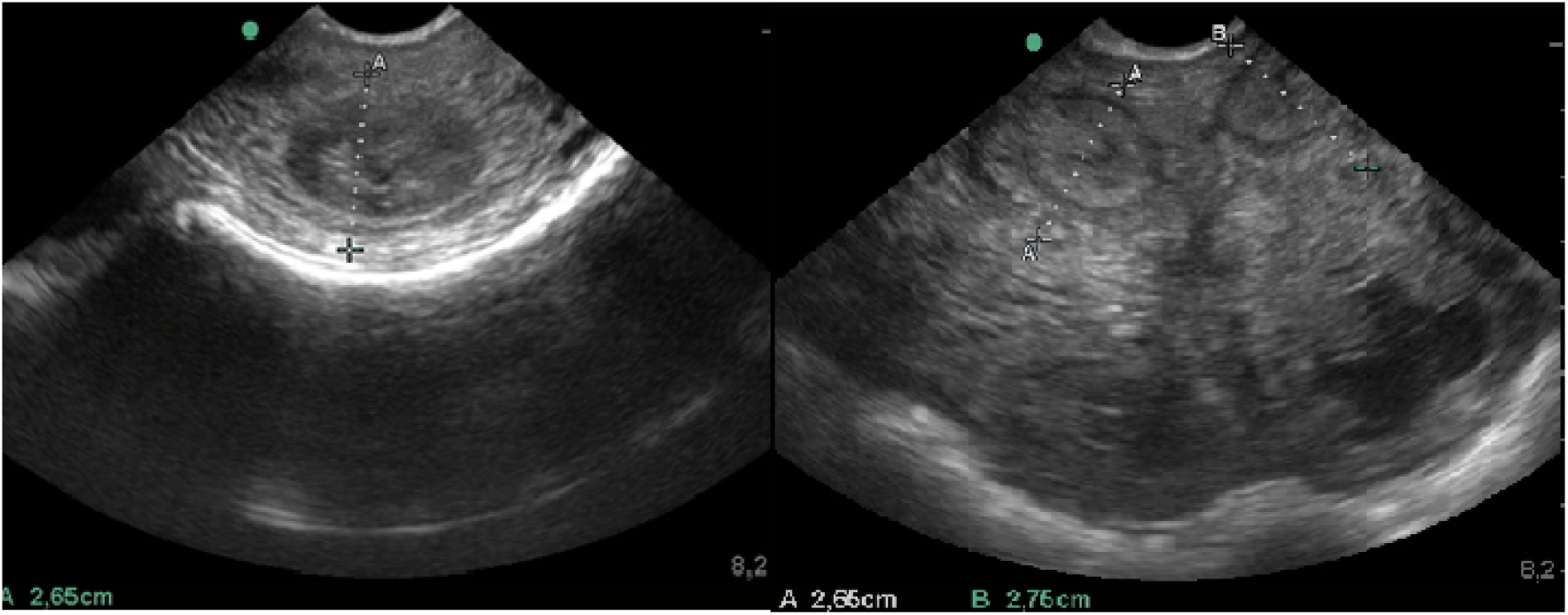
Ultrasound image in transverse section of the cervix (left) and uterine horns (right), using micro probe convex 8.5 Mhz. Measurement in centimeters of the diameter of cervix and uterine horns in bovine. Source: (RIBEIRO, B.L.M. 2016) [21]

**Figure 2 :**
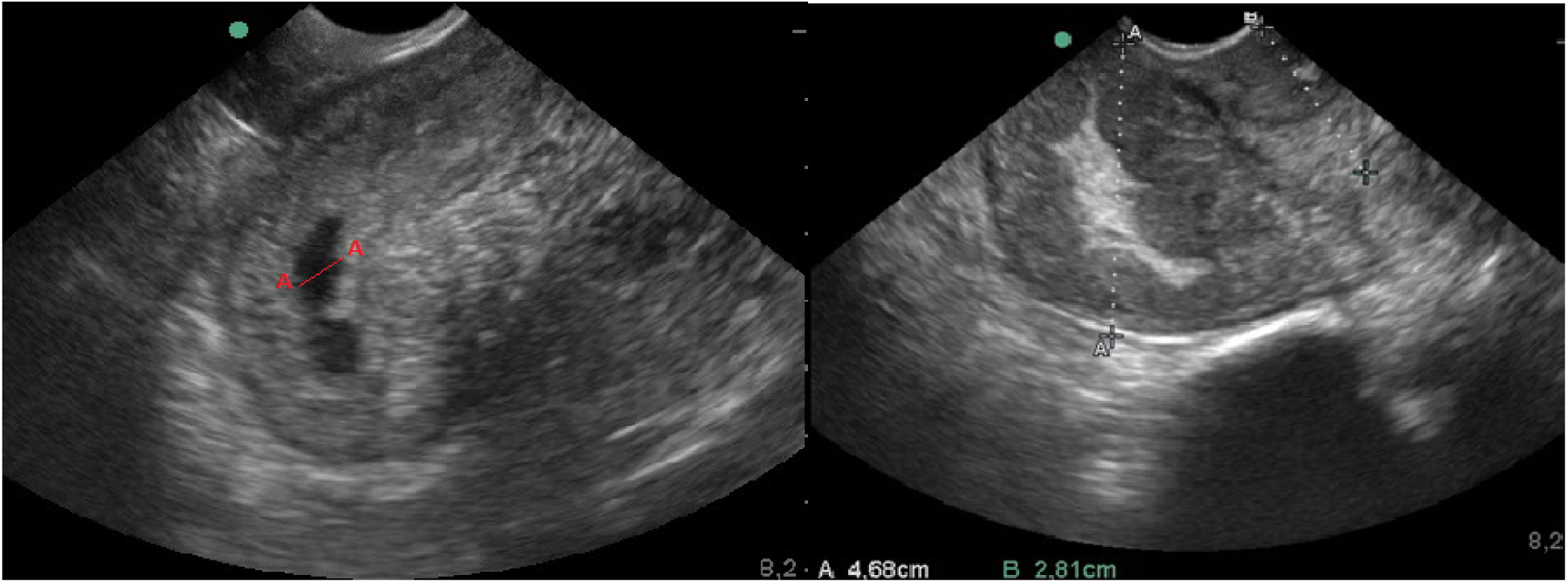
Ultrasound imaging in transverse section of the uterus, using a microconvex 8.5 Mhz probe, characterizing intrauterine contents in abnormal proportions; intrauterine-fluid (left) and hyperechoic intrauterine content. Source: (RIBEIRO, B.L.M. 2016) [21]

After the conventional ultrasonographic evaluation, hemodynamic patterns of the left and right uterine arteries were analyzed using the spectral Doppler mode [12]. Maximum flow velocity, pulsatility index and systole/diastole ratio were determined as described by Heppelmann, Krüger and Leidl (2013) [17] (Figure 3).

**Figure 3:**
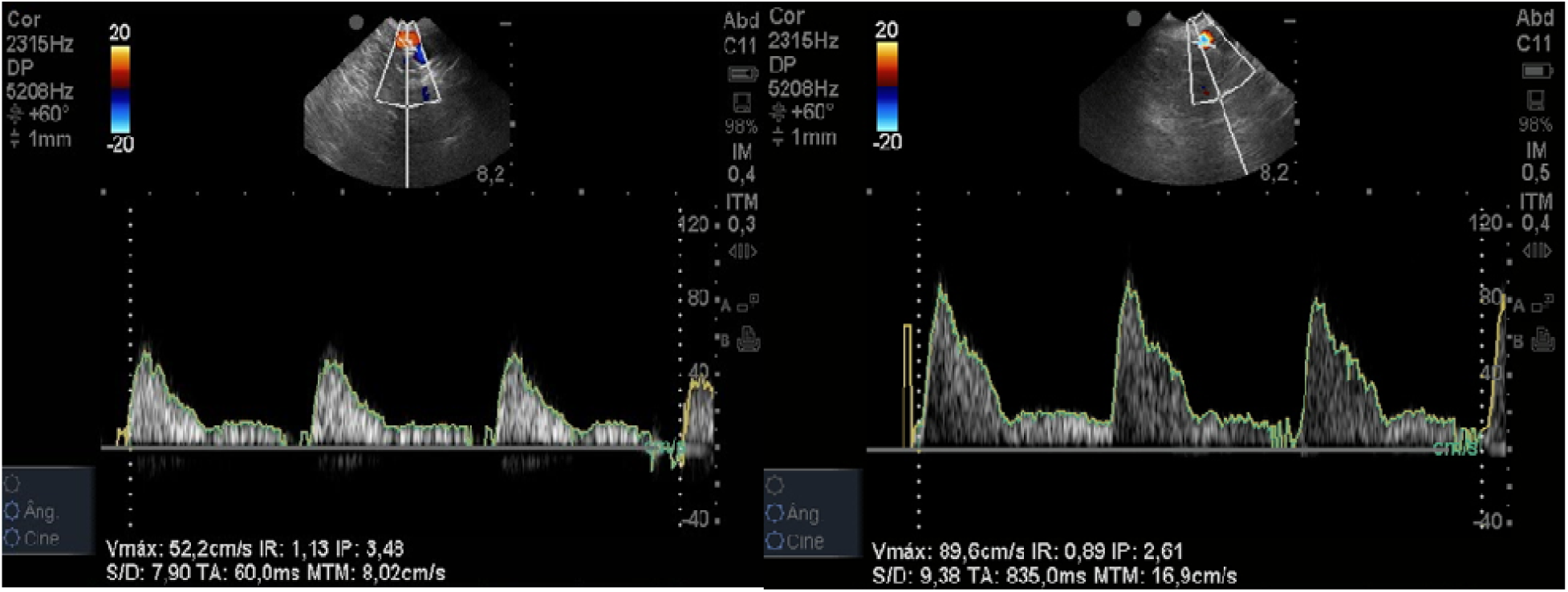
Ultrasound image in cross section of uterus, using 8.5 Mhz micro probe. Spectral Doppler of right left uterine arteries. Source: (RIBEIRO, B.L.M. 2016) [21]

Color Doppler mode was used to evaluate the hemodynamic pattern of the uterine tissue in the intercornual ligament. The colorimetric scale for reproductive patterns [19] was used to establish a parallel with the inflammatory process. The procedures involved a subjective evaluation of the endometrial vascularization (0 -without vascularization to 2 -very vascularized) and the mesometrium (0 -without vascularization to 4 -extremely vascularized), as described by Ginther (2007) [19] (Figures 4 to 8).

**Figure 4:**
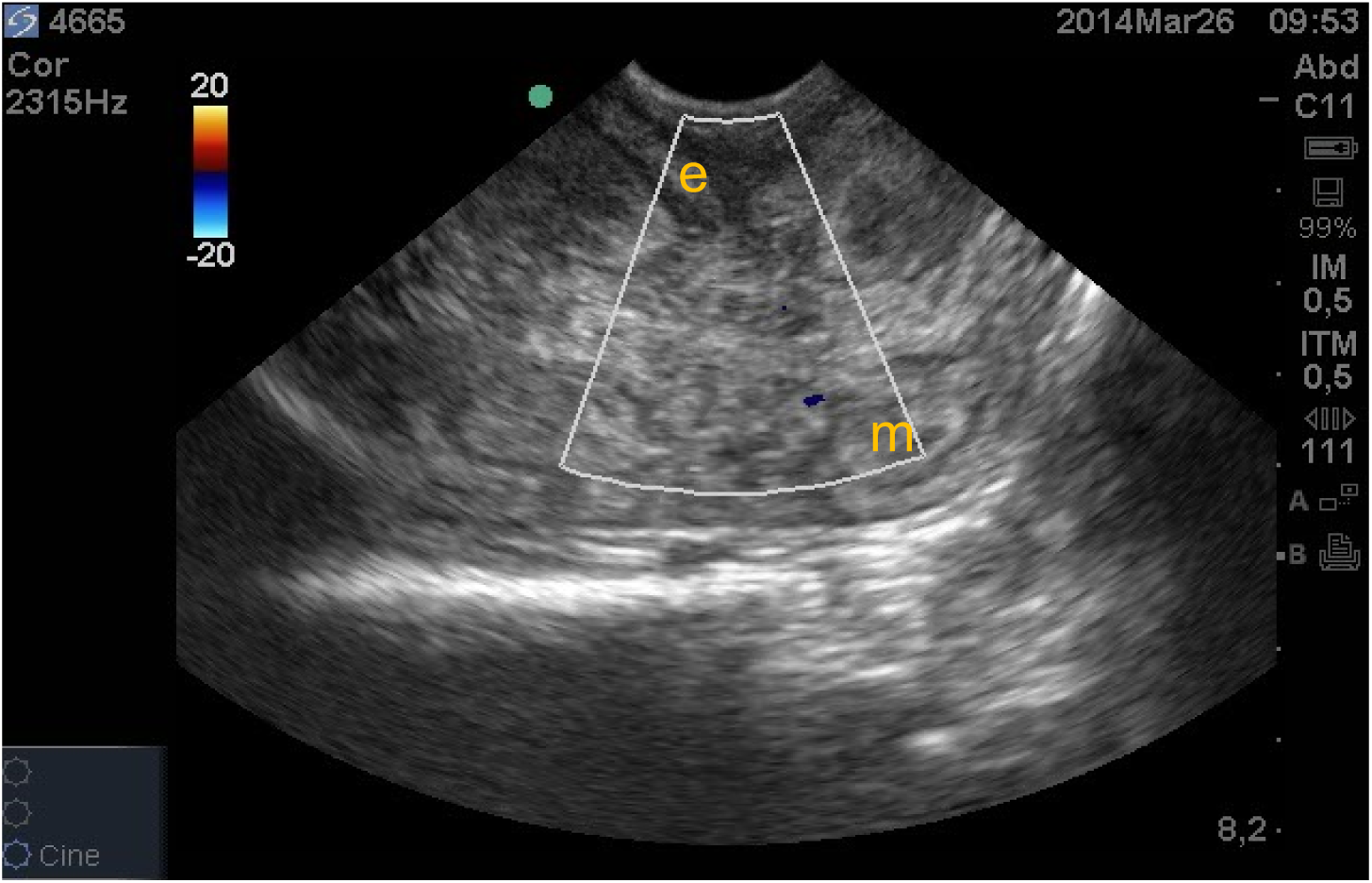
Characterization of 0 score for the subjective evaluation of the vascularization pattern of mesometrium (M) and 0 score for endometrium (e) by colorimetric evaluation technique in Doppler color mode. Source: (RIBEIRO, B.L.M. 2016) [21]

**Figure 5:**
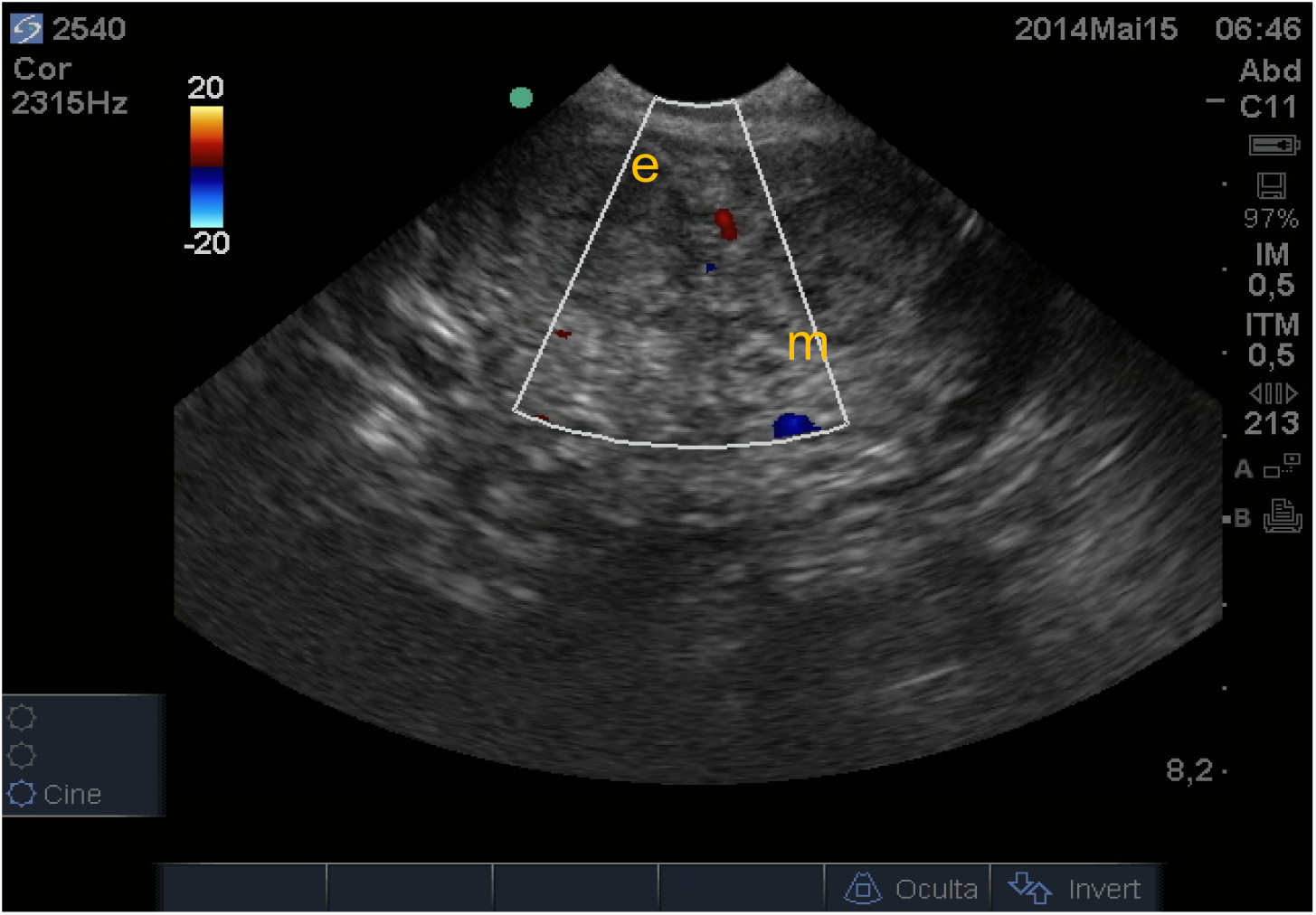
Characterization of a score 1 for the subjective evaluation of the vascularization pattern of mesometrium 0 by colorimetric evaluation technique in Doppler color mode. Source: (RIBEIRO, B.L.M. 2016) [21]

**Figure 6:**
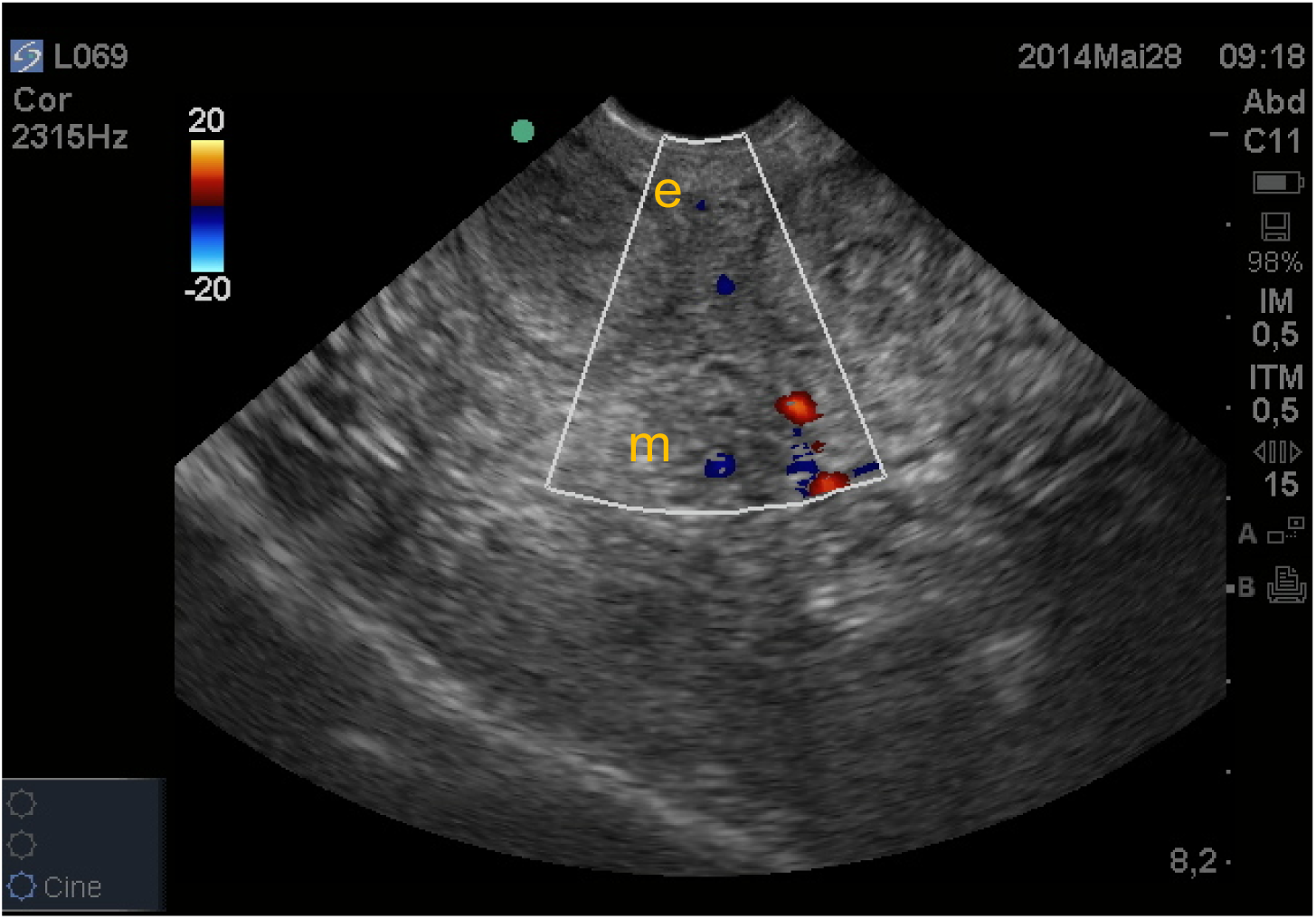
Characterization of score 2 for the subjective evaluation of the pattern of vascularization of the mesometrium and score 0 for endometrium by colorimetric evaluation in Doppler color mode. Source: (RIBEIRO, B.L.M. 2016) [21]

**Figure 7:**
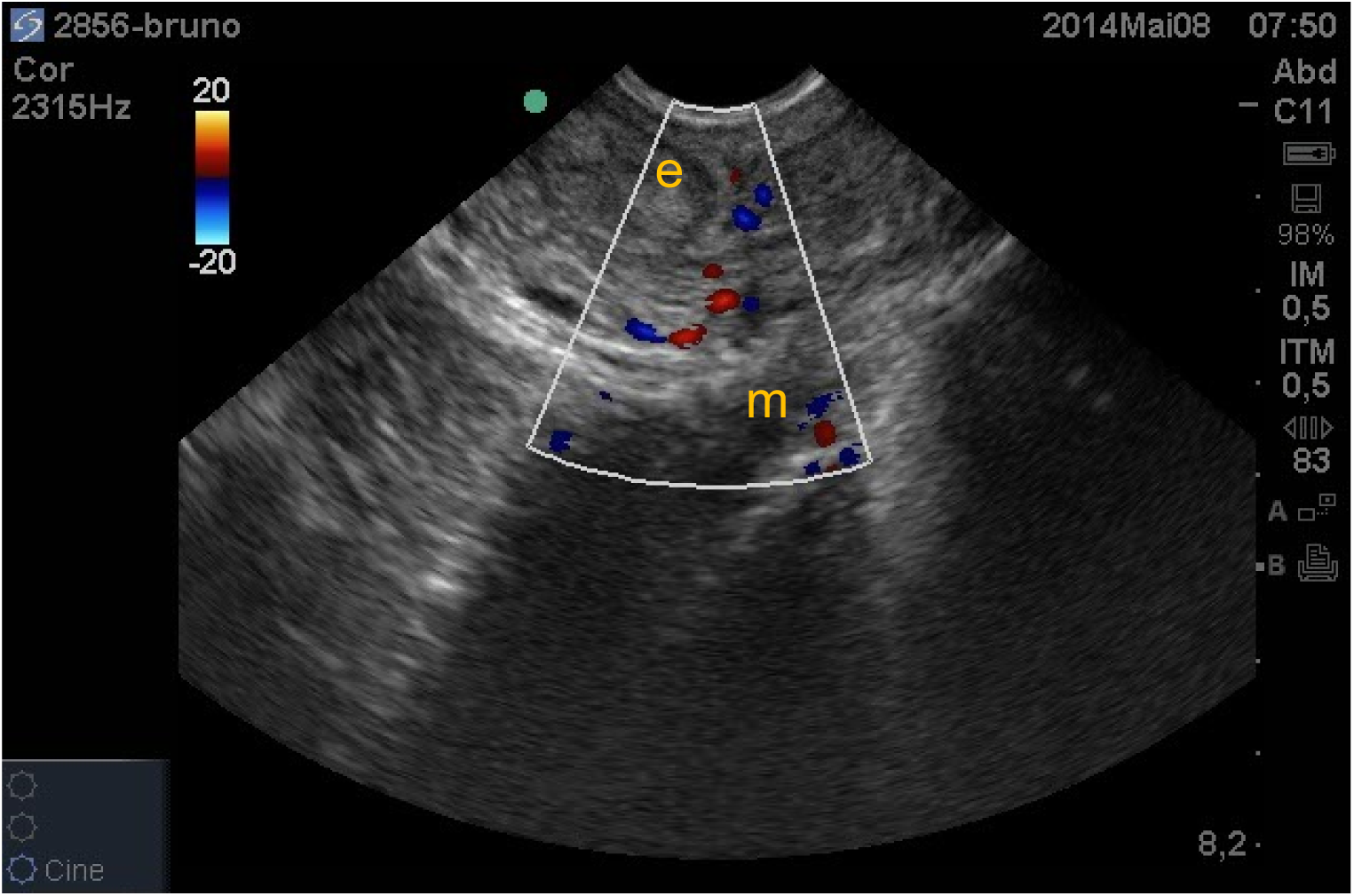
Characterization of score 3 for the subjective evaluation of the vascularization pattern of the mesometrium (m) and score 1 for endometrium (e) by colorimetric evaluation technique in color Doppler mode. Source: (RIBEIRO, B.L.M. 2016) [21]

**Figure 8:**
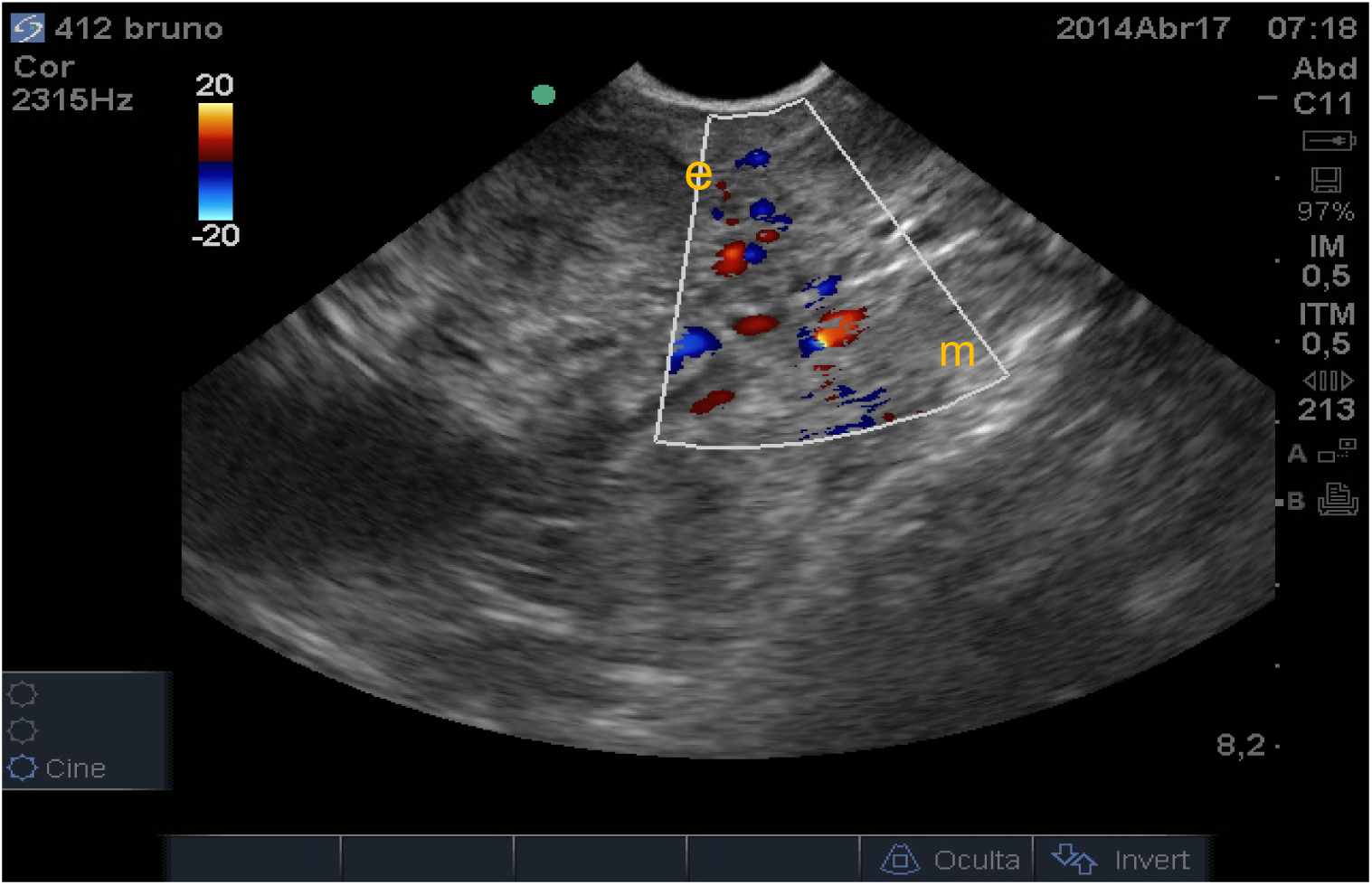
Characterization of the score 4 for the subjective evaluation of the vascularization pattern of the mesometrium (m) and score 2 for the endometrium (e) by colorimetric evaluation in Doppler color mode. Source: (RIBEIRO, B.L.M. 2016) [21]

The ovaries were evaluated for their size, consistency and structures (CL, follicles and cysts). Morphological characteristics of the ovaries were assessed to verify the presence of abnormalities such as cysts or tumors, to determine the existence of cyclic luteal ovarian activity and to estimate the probable phase of the cycle, factors that may affect animal’s fertility [20]. Particularly, the vascularization of the CL was assessed by colorimetric Doppler mode, and it was classified as containing or not containing vascularization (Figure 9).

**Figure 9:**
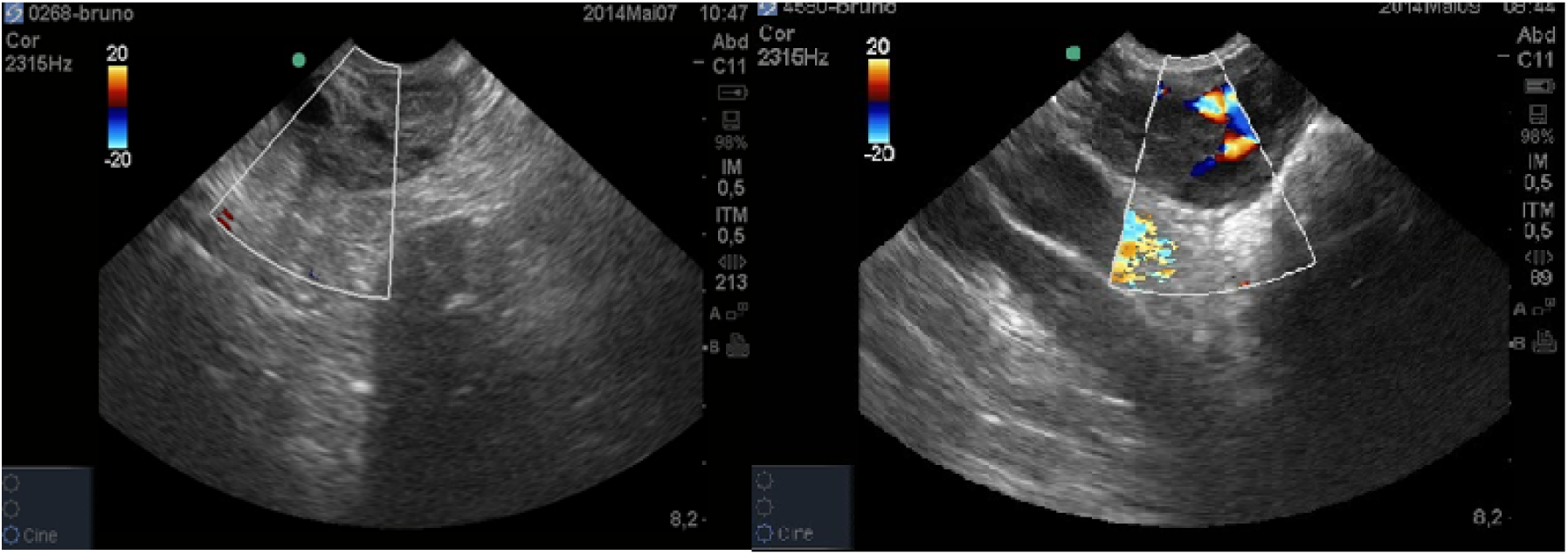
Ovarian evaluation with Doppler color mode: ovary with CL without vascularization (left); ovary with CL vascularized (right). Source: (RIBEIRO, B.L.M. 2016) [21]

### 2.5 Hysteroscopy

The hysteroscopic examination was performed using a rigid endoscope (430 × 6 mm) with 2 channels: one for the passage of 0.9% sterile saline for uterine lumen distension and another for cytological and microbiological samples collection. The optic part of the endoscope was introduced into the vagina, guided by transrectal palpation. Once inside the uterus, the sterile saline was infused to explore this anatomic site. The presence of mucopurulent or purulent inflammatory lesions in the mucosa were considered as signs of endometritis [22]. The images were classified as healthy or unhealthy (presence of pus, fibrin and hyperemia) mucosa. After exploration, a small portion of saline was recovered and used for macroscopic, cytological and microbiological evaluation of the uterus content. After each examination, the optic cable was washed with water only. The work channels were brushed with a flexible 75cm X 0.20mm brush immersed in enzymatic detergent (Hs-enzyme®, Strattner®) and, then, disinfected using paracetic acid (Peroxylife®), according to the manufacturer’s instructions.

### 2.6 Uterine cytology

One hundred and fifty microliters of the recovered saline were transferred to a cytocentrifuge chamber and samples were fixed and centrifuged on a glass slide for microscopy at approximately 550 rpm for 6 minutes [6]. The slides were stained using the Panotipo Rapido® kit, and cytology was performed by counting 100 cells, under magnification of 400x using an optical microscope. The percentage of neutrophil polymorphonuclear cells (% pmn) (Figure 12) was determined. Cows with %Ne > 10% were considered positive to the cytology test [23].

### 2.7 Microbiological

Five hundred microliters of the recovered saline were added to thioglycolate broth (Difco®) and kept at 2°-10 °C until processing at the Laboratory of General Bacteriology, Biological Institute, São Paulo. Ten microliters were plated in 5% sheep blood agar and incubated for 72 hours at 37 °C under microaerophilic conditions. Morphological characteristics of each colony and Gram staining were performed. Bacteria species were determined using biochemical tests according to Winn et al. (2008).

### 2.8 Statistical analysis

Error normality and homogeneity of variance were analyzed by Shapiro-Wilk Test and Bartlett Test, respectively. Non-normal data were analyzed using a non-parametric test (Kruskal-Wallis test) (proc npar1way). Differences between the groups (control and endometritis) were evaluated by ANOVA, reaffirmed by a t-test (JMP 12.0 -SAS). Same conditions (proc glimmix from SAS) were used for binary data (“subjective evaluation score of endometrial perfusion” and “subjective evaluation score of mesometrium perfusion”). Correlation between variables were performed using Spearmann correlation test for variables that did not present normal distribution (proc corr from SAS). A significance level of 5% was used for all tests. All tests were performed in SAS.

## 3. Results

One hundred cows were initially examined, but due to problems with the ultrasound device, eleven samples were lost. So, 89 cows were enrolled in this study, and they were divided accordingly to citology results into two groups: control group (E) (N= 33) and endometritis group (N = 56).

### 3.1 Ultrasonographic exam

Cows with endometritis had the cervix (*P* = 0.040) and the left horn (*P* = 0.020) increased compared to healthy cows (Table 1). In the endometritis group, 78.6% of the cows showed abnormal uterine discharge, while in health group only 57.6% of the animals showed this same condition (*P =* 0.0005) (Table 2). Besides, intrauterine heterogeneous content were increased in sick cows (66.0%) compared to healthy cows (30.3%) (*P* = 0.0011) (Table 3). Evaluation of endometrial and mesometrium vascularization by the colorimetric method revealed differences in the vascularization score (vc) of the endometrium between healthy (C) and endometritis group (C) (*P <* 0.05) (Table 4 and 5). During ovarian examination, presence or absence of CL were not diferent between group (C) and group (E) (Table 6). In addition, the absence of vascularizated corpus luteum in the left ovary was increased in the endometritis group (*P* = 0.029) (Table 7). The spectral Doppler evaluation of the uterine arteries revealed no differences between groups (Table 8).

**Table 1.** Average diameter between measurements of linear ultrasonography of the cervix and uterine horns with Endometritis

| Uterine structures | Control | Endometritis | P value |
| --- | --- | --- | --- |
| Cervical (cm) | 3.55($\pm 0.13$) | 3.89($\pm 0.01$) | 0.040 |
| Right horn (cm) | 2.56($\pm 0.11$) | 2.77($\pm 0.08$) | 0.100 |
| Left horn (cm) | 2.48($\pm 0.13$) | 2.89($\pm 0.10$) | 0.020 |

Source: (RIBEIRO, B.L.M., 2016).

**Table 2.**
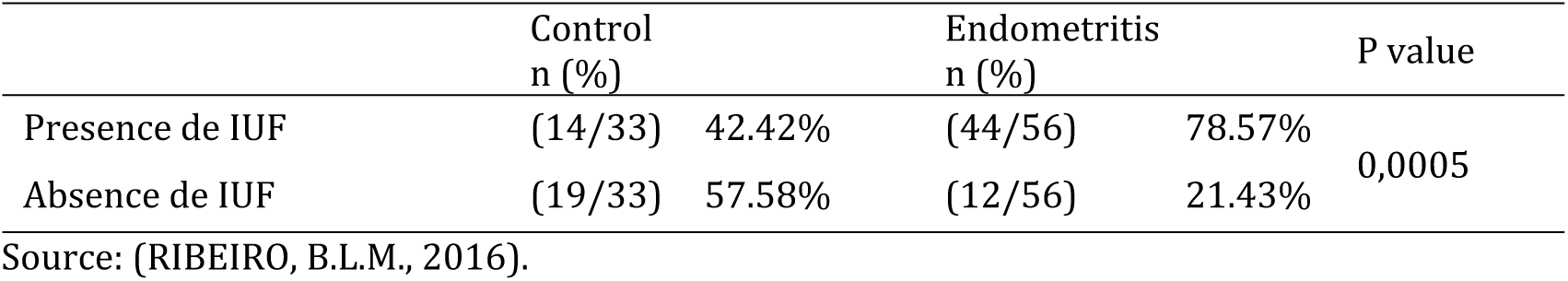
Presence of intrauterine fluid (IUF) in cows with and without Endometritis.

|  | Control<br>n (%) | Endometritis<br>n (%) | P value |
| --- | --- | --- | --- |
| Presence de IUF | (14/33) 42.42% | (44/56) 78.57% | 0,0005 |
| Absence de IUF | (19/33) 57.58% | (12/56) 21.43% |  |

Source: (RIBEIRO, B.L.M., 2016).

**Table 3.** Animals that presented intrauterine heterogeneous content (IUHC) in abnormal amounts throughout the experiment

|  | Control<br>n (%) |  | Endometritis<br>n (%) |  | P value |
| --- | --- | --- | --- | --- | --- |
| Presence de IUHC | (10/33) | 30.30% | (37/56) | 66.07% | 0,0011 |
| Absence de IUHC | (23/33) | 69.70% | (19/56) | 33.93% |  |

Source: (RIBEIRO, B.L.M., 2016).

**Table 4.**
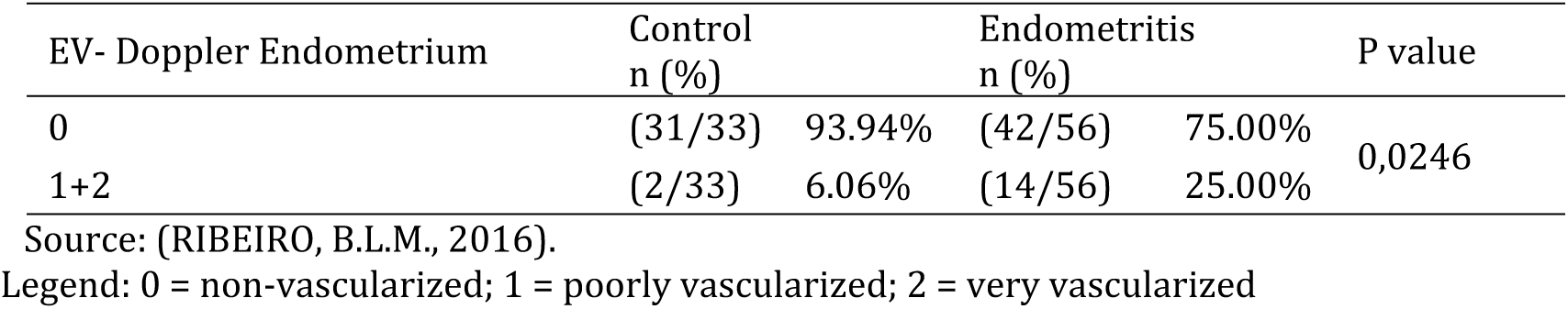
Evaluation of vascular pattern of endometrium.

**Table 5.** Frequency of evaluation of the vascular pattern of Mesometrium.

| EV- Doppler Mesometrium | Control<br>n (%) |  | Endometritis<br>n (%) |  | P value |
| --- | --- | --- | --- | --- | --- |
| 0 | (10/33) | 30.30% | (8/56) | 14.29% | 0,0344 |
| 1 | (11/33) | 33.33% | (9/56) | 16.07% |  |
| 2 | (5/33) | 15.15% | (14/56) | 25.00% |  |
| 3 | (5/33) | 15.15% | (12/56) | 21.43% |  |
| 4 | (2/33) | 6.06% | (13/56) | 23.21% |  |

Source: (RIBEIRO, B.L.M., 2016).
Legend: 0 = non-vascularized; 1 = poorly vascularized; 2 = vascularized; 3 = very vascularized; 4 = extremely vascularized.

**Table 6.** Frequency of Presence of CL as a function of Endometritis.

|  | Control<br>n (%) |  | Endometritis<br>n (%) |  | P value |
| --- | --- | --- | --- | --- | --- |
| Presence de CL Right | (15/33) | 45.45% | (24/56) | 42.86% | 0.811 |
| Absence de CL Right | (18/33) | 54.55% | (32/56) | 57.14% |  |
| Presence de CL Left | (16/33) | 48.48% | (18/56) | 32.14% | 0.125 |
| Absence de CL Left | (17/33) | 51.52% | (38/56) | 67.86% |  |

Source: (RIBEIRO, B.L.M., 2016).

**Table 7.** Presence of CLv as a function of Endometritis.

|  | Control<br>n (%) |  | Endometritis<br>n (%) |  | P value |
| --- | --- | --- | --- | --- | --- |
| Presence de CLv Right | (12/33) | 36.36% | (19/56) | 33.93% | 0.815 |
| Absence de CLv Right | (21/33) | 63.64% | (37/56) | 66.07% |  |
| Presence de CLv Left | (15/33) | 45.45% | (13/56) | 23.21% | 0.029 |
| Absence de CLv Left | (18/33) | 54.55% | (43/56) | 76.79% |  |

Source: (RIBEIRO, B.L.M., 2016).

**Table 8.**
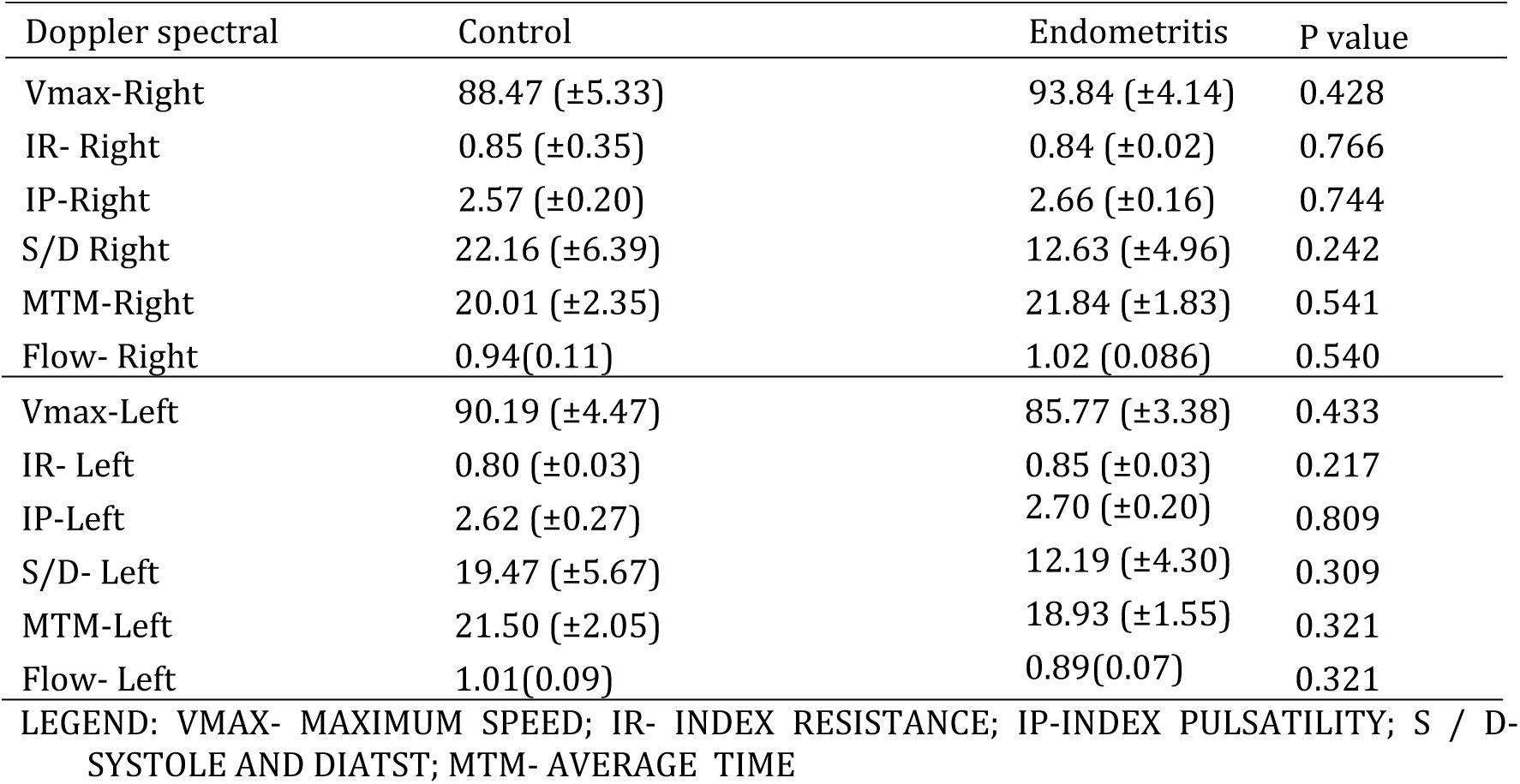
Spectral Doppler average in Uterine Arteries.

| Doppler spectral | Control | Endometritis | P value |
| --- | --- | --- | --- |
| Vmax-Right | 88.47 (±5.33) | 93.84 (±4.14) | 0.428 |
| IR- Right | 0.85 (±0.35) | 0.84 (±0.02) | 0.766 |
| IP-Right | 2.57 (±0.20) | 2.66 (±0.16) | 0.744 |
| S/D Right | 22.16 (±6.39) | 12.63 (±4.96) | 0.242 |
| MTM-Right | 20.01 (±2.35) | 21.84 (±1.83) | 0.541 |
| Flow- Right | 0.94(0.11) | 1.02 (0.086) | 0.540 |
| Vmax-Left | 90.19 (±4.47) | 85.77 (±3.38) | 0.433 |
| IR- Left | 0.80 (±0.03) | 0.85 (±0.03) | 0.217 |
| IP-Left | 2.62 (±0.27) | 2.70 (±0.20) | 0.809 |
| S/D- Left | 19.47 (±5.67) | 12.19 (±4.30) | 0.309 |
| MTM-Left | 21.50 (±2.05) | 18.93 (±1.55) | 0.321 |
| Flow- Left | 1.01(0.09) | 0.89(0.07) | 0.321 |

LEGEND: VMAX- MAXIMUM SPEED; IR- INDEX RESISTANCE; IP-INDEX PULSATILITY; S / D- SYSTOLE AND DIATST; MTM- AVERAGE TIME

### 3.3 Microbiological findings

*Bacillus spp., Trueperella pyogenes, Escherichia coli* and *Staphylococcus intermedius* were the most isolated bacteria among samples. Yeast was also detected in 25% of samples (Table 9).

**Table 9.** Identification of the microorganisms in samples.

|  | Endometritis |  |
| --- | --- | --- |
|  | n (%) |  |
| Bacterium+Yeast | (6/85) | 7.06% |
| Yeast | (21/85) | 24.71% |
| Bacterium | (38/85) | 44.71% |
| *T. pyogenes* | (10/85) | 11.76% |
| *S. intermedius* | (7/85) | 8.24% |
| *S. auerus* | (1/85) | 1.18% |
| *Streptococcus* sp. | (1/85) | 1.18% |
| *Bacillus* sp.; | (13/85) | 15.29% |
| *E. coli* | (7/85) | 8.24% |
| *Proteus* sp. | (2/85) | 3.53% |

Source: (RIBEIRO, B.L.M., 2016).

### 3.4 Correlations with variables

A positive correlation between mixed culture (bacteria + yeasts) and endometritis was observed (*P* = 0.0003). In addition, weak correlations were observed between intrauterine heterogeneous content (r = 0.34), intrauterine fluid (r = 0.37) and endometritis. The disease was also correlated with Doppler ultrasonography (*P* = 0.0027) and endometrial Doppler (*P* = 0.0383). A weak and negative correlation with the presence of corpus luteum in the vascularized left ovary (*P* = 0.0359) and endometritis was also detected (Table 10).

**Table 10.**
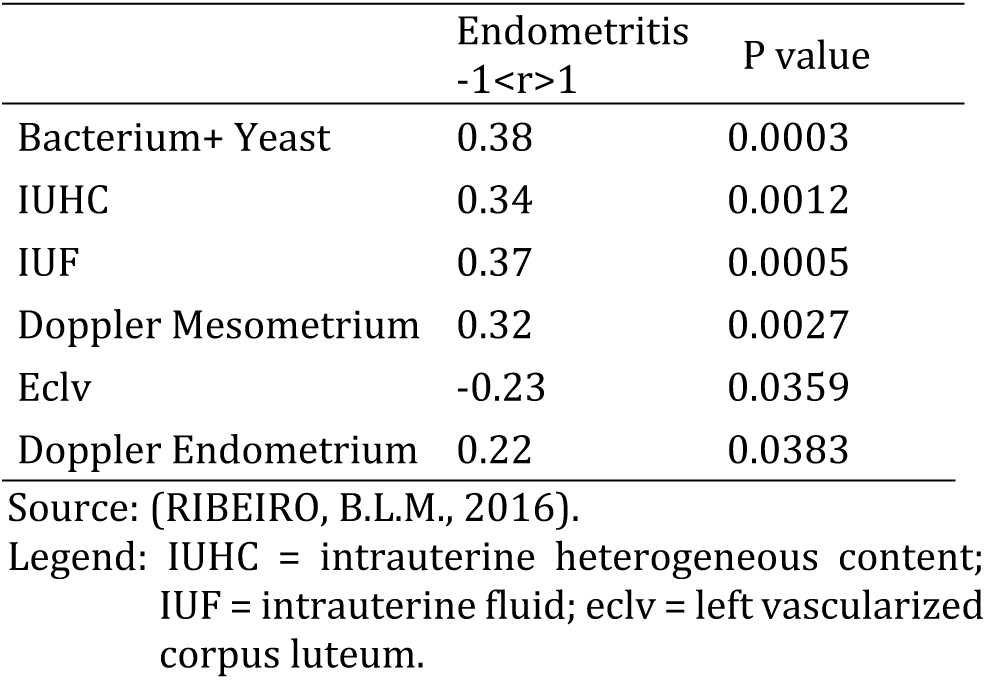
Correlation between Endometritis and other variables

Intrauterine fluid (IUF) was correlated with *Trueperella pyogenes* (r = 0.27), yeast (r = 0.25), *Staphylococcus intermedius* (r = 0.23), and *Escherichia coli* (r = 0.23) (*P* < 0.05) (Table 13). Strong correlations between intrauterine heterogeneous content (IUHC), and IUF (*P = 0.0001*) and the presence of bacteria (P = 0.0001) were detected. IUHC was also correlated with *Trueperella pyogenes* (r = 0.36), *Escherichia coli* (r = 0.30), *Bacillus spp.* (r = 0.25), *Staphylococcus intermedius* (r = 0.22) and yeast (*P* < 0.05) (Table 14). Endometrial Doppler was correlated with *Trueperella pyogenes* (*P =* 0.0003) and IUHC (*P* = 0.0047) (Table 15). Finally, mesometrial Doppler was correlated with endometrial Doppler (*P* < 0.001), uterine bacteria (*P =* 0.001) and IUHC (*P =* 0.049). A negative correlation with the presence of yeast (*P* = 0.022) was also observed (Table 16).

**Table 11.**
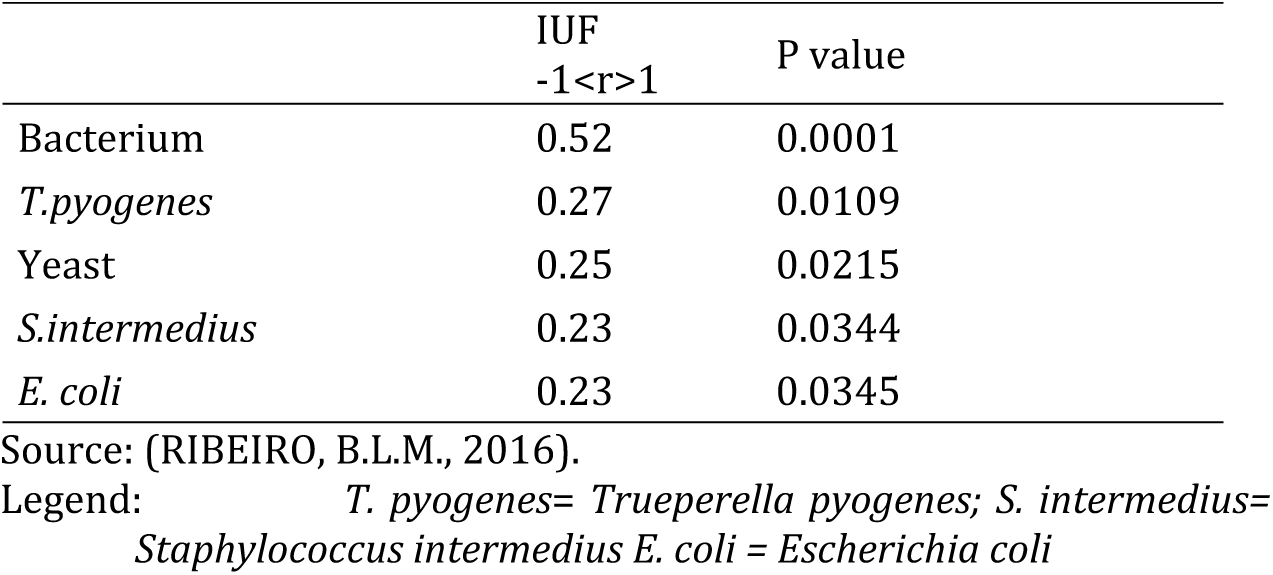
Correlation between Presence of Intrauterine Fluid (IUF) compared to Presence of microorganisms

**Table 12.** Correlation of animals with heterogeneous content compared to other variables.

|  | IUHC<br>-1<r>1 | P value |
| --- | --- | --- |
| IUF | 0.72 | 0.0001 |
| Bacterium | 0.64 | 0.0001 |
| *T. pyogenes* | 0.36 | 0.0007 |
| *E. coli* | 0.3 | 0.0052 |
| Yeast | 0.27 | 0.0135 |
| *Bacillus* sp. | 0.25 | 0.0223 |
| *S. intermedius* | 0.22 | 0.0434 |

Source: (RIBEIRO, B.L.M., 2016).
Legend: IUF = Intrauterine fluid; *T. pyogenes* = *Trueperella pyogenes*; *S. intermedius*= *Staphylococcus intermedius*; *E. coli* = *Escherichia coli*.

**Table 13.** Endometrium color Doppler correlation compared to other variables.

|  | Doppler Endometrium<br>-1<r>1 | P value |
| --- | --- | --- |
| ***T. PYOGENES*** | 0.38 | 0.0003 |
| Bacterium | 0.38 | 0.0003 |
| IUHC | 0.36 | 0.0047 |

Source: (RIBEIRO, B.L.M., 2016).
Legend: IUHC = intrauterine heterogeneous content; *T. pyogenes* = *Trueperella pyogenes*.

**Table 14.** Doppler color correlation of Mesometrium compared with other variables

|  | Doppler<br>Mesometrium | P value |
| --- | --- | --- |
| Doppler<br>Endometrium | 0.52 | 0.000 |
| Bacterium | 0.35 | 0.001 |
| Yeast | -0.25 | 0.022 |
| IUHC | 0.21 | 0.049 |

Source: (RIBEIRO, B.L.M., 2016).
Legend: IUHC = intrauterine heterogeneous content.

## 4. Discussion

Endometritis is a uterine disease and is related to negative reproductive performance [8]. Besides physical examination, new diagnostic tools are important to improve the precocity in the diagnosis of this disease [24].

Our results showed differences between the size of the cervix and a larger uterus in cows with endometritis. After delivery, the uterus is greatly enlarged (approximately 8 to 10 kg). In general, the uterine macroscopic involution occurs between 3 to 5 weeks postpartum, when the uterus should weight approximately 0.9 kg and the diameter of the pre-gravid uterine horn should be less than 5 cm, total involution of the cervix occurs between 4 to 6 weeks postpartum [25]. A new pregnancy depends on the anatomical and functional return of the genital tract to a state similar to that before pregnancy [26]. In the present research, the increased left horn in animals with endometritis was observed. This information is in agreement with other researches that proved that a uterine infection provides a delayed involution [27, 26]. In this study, cows with endometritis presented larger cervix diameter compared to healthy animals. LeBlanc et al. (2002) [5] highlighted that cows with clinical endometritis showed cervical measurement ≥7.5 cm in diameter after 20 days postpartum and they presented mucopurulent or purulent discharge detected by vaginoscopy after 26 days postpartum. Our study highlights that endometritis was increased in cows showing abnormal intrauterine fluid accumulation, and it was in agreement with several studies [28, 29, 16]. Intrauterine fluid was correlated with *Trueperella pyogenes*, yeast, *Staphylococcus intermedius, Escherichia coli*, which suggests that these agents cause inflammation of the uterine mucosa and proliferation of the endometrial glands, leading to a greater accumulation of intrauterine fluid.

Endometritis was highly detected in cows showing intrauterine heterogeneous content (IHC). Meira Jr. et al. (2012) [16] also described high correlation between IHC and the diagnosis of endometritis. IHC was also associated with *Trueperella pyogenes, Escherichia coli, Bacillus spp*. and *Staphylococcus intermedius*, and yeast, demonstrating that these microorganisms may cause genital discharge [16].

The presence of vascularization in the endometrium (degrees 1 and 2) was increased in cows with endometritis, suggesting differences between healthy and unhealthy cows. The color Doppler mode provides colored images of the blood fow, allowing estimation of the tissue vascularization [34]. Ginther (2007) [19] proposed a model to evaluate the uterine hemodynamic associated with endometritis. This model was modified in the present research, and our results showed a few animals with alterations in the subjective evaluation score of endometrial vascularization, and a positive correlation with *Trueperella pyogenes*.

Ultrasonographic evaluation of the endometrium using color Doppler mode showed a positive correlation with endometritis. This information emphasizes the possible combination of this new noninvasive technique with the standard diagnostic methods to predict uterine disease. No difference was detected at data analisys of spectral Doppler mode. Herzog and Bollwein (2013) [2] reported increased pulsatility at 24 hours postpartum with a peak on the 28th day, decreasing progressively until 90th day. An inverse proportional relation was observed in blood flow, which already starts high in the postpartum and decreases until around the 28th day maintaining basal levels up to 90th day. Clinical and histologic uterine involution ranges from 21 to 50 postpartum, and this may differs from the recovery of hemodynamic patterns of the uterus that requires more time [30].

Advancing the understanding of uterine perfusion, Bollwein et al. (2002) [13] when evaluating pregnant animals, concluded that throughout pregnancy there is a strong increase of blood flow in the uterine arteries. However, they observed that the resistivity index decreased in the first 8 months of gestation and remained at a relatively constant level until the calf was born. It is known that during the diestro the blood flow velocity remains constant and are correlated to the plasma concentration estrogen and progesterone, indicating a moderate positive correlation Bollwein et al. (2016) [30]. These results indicate that there are other factors involved in the regulation of blood flow [31].

In healthy cows the uterine blood flow decreased after the complete uterine involution. However, cows with uterine disease presents slow uterine involution, occurring between 45 and 65 days postpartum. These results indicate an association between incomplete uterine involution and regeneration of the uterine vascular layer in cows with puerperal disease. Changes in uterine perfusion are greatly pronounced during the first 4 days after delivery. Uterine puerperal diseases such as retention fetal attachments and metritis have a negative impact on uterine involution [17]. Thus, uterine blood flow is affected by puerperal uterine diseases. Cows with retained placenta have higher resistivity index than healthy cows. On the 8th day postpartum, cows with metritis had higher blood flow and decreased pulsatility in the arteries compared to healthy cows [17].

Evaluation of the uterine and ovarian vascularization helps to determine the best moment to perform artificial insemination [31]. The vascular perfusion of corpus luteum and uterus is greater during the first follicular wave compared to the second follicular wave of the estrous cycle [31]. The color Doppler mode identifies follicles in normal development and predicts the next ovulate. In this research, the ovarian evaluation showed that some cows with endometritis had no corpus luteum in both the right and left ovary. However, animals with right and left CL presented 61.54% and 52.94% vascularized, respectively. Color Doppler mode showed that cows with no vascularized left and right corpus luteum had endometritis, demonstrating that there is a interference in reproduction performance even in endometritic cows exhibiting normal estrous cycle.

Microbiological results showed the presence of *Bacillus* spp., *Trueperella pyogenes, Staphylococcus intermedius, Escherichia coli, Streptococcus* spp., *Proteus* spp., *Staphylococcus aureus, Enterobacteria and Serratia* spp. besides fungus and yeast. Santos, Gilbert and Bicalho (2011) [32] reported that *Escherichia coli, Streptococcus* spp., *Trueperella pyogenes* and *Fusobacterium necrophorum* are the main bacteria that commonly contaminate the uterine lumen after childbirth. These microorganisms were also associated with some uterine diseases. *Trueperella pyogenes* acts in synergisms with *Fusobacterium necrophorum, Bacteroides* spp., and *Prevotella* spp. [33]. Potter et al. (2010) [34] reported that *Trueperella pyogenes* is recognized as an important pathogen associated with endometritis due to its persistence in the contaminated uterus. Recently, Boer et al. (2015) [35] found that cows with any bacterial growth at 21 days postpartum, regardless of bacterial species, had a lower conception rate. Using the 16S rRNA sequencing, Machado et al. (2012) [36] detected *Trueperella pyogenes* increased in cows with endometritis. Yeasts have been associated with chronic endometritis in women [40]. In the present research, yeasts were detected in 22.5% of cows. The onset of *Trueperella pyogenes* was correlated with uterine inflammation seen by the colorimetric ultrasonographic method.

Although endometritis has a spontaneously healing condition, Dubuc et al. (2011) [7] found 63% of spontaneous cure, while LeBlanc et al. (2002a) [5] described 77% self-healing. It still causes many economic and reproductive damages in the dairy industry. In order to reduce this problem, it is important to use new tools to improve the precision and the precocity in the diagnostic, upgrading the reproductive indices.

## 5. Conclusions

It is detected the relationship between the diameter of the cervix and the uterine horns with endometritis. This relationship was notorious when the presence of intrauterine fluid and intrauterine heterogeneous content was verified. Color Doppler showed the association between the vascularization of inflamed tissue and endometritis. However, performing spectral Doppler ultrasound showed no differences between healthy and unhealthy animals. The microbiological examination showed that *Trueperella pyogenesis* and *Escherichia coli* have an important role in the development of endometritis, corroborating other researches. Less invasive techniques with fast results such as Doppler ultrasonography can provide satisfactory answers regarding to the evolution of uterine alterations, improving reproductive rates.

## Acknowledgements and Funding

Authors thank São Paulo Research Foundation (Process number 2014/02676-4)

